# Amylo-Pipe: an integrated web server for mechanistic and kinetic prediction of protein and peptide aggregation

**DOI:** 10.64898/2026.06.09.731090

**Authors:** Puneet Rawat, R. Prabakaran, Niccolò Cardente, Sandeep Kumar, Victor Greiff, M. Michael Gromiha

## Abstract

Protein aggregation is central to amyloid-related disorders and remains a major developability challenge for protein therapeutics. Over the past two decades, significant advances have been made to predict aggregation-prone regions (APRs) and estimate aggregation propensity in proteins and peptides. In contrast, the prediction of aggregation kinetics has received relatively less attention due to the limited availability and heterogeneity of experimental data. Consequently, aggregation propensities from APR prediction algorithms were widely accepted as a means to predict relative changes in the aggregation kinetics of proteins and mutants. Previous studies have demonstrated, using large-scale datasets, that aggregation propensity shows a weak or inconsistent correlation with aggregation kinetics. In the present study, we have integrated complementary state-of-the-art mechanistic and kinetic prediction tools for protein aggregation into a unified, user-friendly web framework entitled “Amylo-Pipe”. Amylo-Pipe also implements practical features that are especially useful for protein engineering, such as gatekeeper-residue mutational scanning to support the design of aggregation-resistant variants. By consolidating multiple prediction tasks in a single interface, Amylo-Pipe enables a more comprehensive assessment of aggregation behavior than APR-only workflows. The web server is freely accessible at: https://web.iitm.ac.in/bioinfo2/amylopipe/.

## 1. Introduction

Accumulation of misfolded or unfolded proteins, widely known as “protein aggregation”, is a major cause of several disease conditions in humans (Alzheimer’s disease, Parkinson’s disease, AL amyloidosis, etc.) and a hindrance in development of protein-based products (enzymes, antibody therapeutics, etc.) ^1–3^. It is postulated that almost all proteins can aggregate under appropriate environmental conditions ^4^. Short sequence motifs, called aggregation-prone regions (APRs), that can trigger aggregation in peptides and proteins ^5^. A computational analysis of the human proteome showed that almost three-fourths of the human proteins contain at least one APR ^6^. However, APR detection alone does not provide a complete description of aggregation behavior of a protein or peptide. The presence of charged residues and proline (gatekeeper residues) at the flanks of the APR is well established to hinder the aggregation process ^7–10^. Moreover, the rate at which an APR drives aggregation depends not only on its intrinsic propensity but also on additional factors such as structural context, solvent exposure, conformational stability, local flexibility, and environmental conditions including pH, temperature, ionic strength, and protein concentration ^11–14^. Recent methodological work continues to support the view that APR identification (mechanistic predictions) and aggregation kinetics predictions should be treated as distinct but complementary prediction tasks ^14,15^. Mechanistic prediction primarily estimates the intrinsic aggregation propensity of the proteins and peptides, which leads to identification of APRs. Some of the tools for aggregation propensity include Cordax ^16^, CANYA ^17^, Waltz ^18^, Tango ^5^, ANuPP ^19^ among others. Generally, the APRs are buried into the core of the protein. Therefore, APR prediction also includes consideration of conformational stability and relative solvent accessibility of the APR ^20–24^. Secondly, aggregation kinetics refers to the time-dependent accumulation of protein aggregates under defined environmental conditions ^25^. In contrast to the APR predictions, only a few attempts have been made to predict aggregation kinetics, owing to the limited availability of experimental datasets ^26–29^.

In this work, we have combined the state-of-the-art methods to predict the mechanistic and kinetic aspects of protein aggregation into a platform to provide a detailed overview of the aggregation capability of proteins and peptides. This platform “Amylo-Pipe” includes two mechanistic prediction methods: ANuPP (APR and aggregation propensity prediction) ^19^ and V_L_AmY-Pred (amyloidogenic/non-amyloidogenic antibody light chain prediction) ^30^; two sequence-based kinetics prediction methods AbsoluRATE (quantitative prediction of aggregation rate for protein and peptide) ^25^ and AggreRATE-Disc (classification method to predict aggregation rate enhancer/mitigator point mutation) ^31^; and a structure-based kinetics prediction method AggreRATE-Pred (change in aggregation rate upon point mutation) (**Fig. 1**) ^32^. The web server also contains unique features for comprehensive analysis of a single protein or peptide, such as gatekeeper mutation scanning. The web server can be accessed at https://web.iitm.ac.in/bioinfo2/amylopipe/.

**Figure 1.**
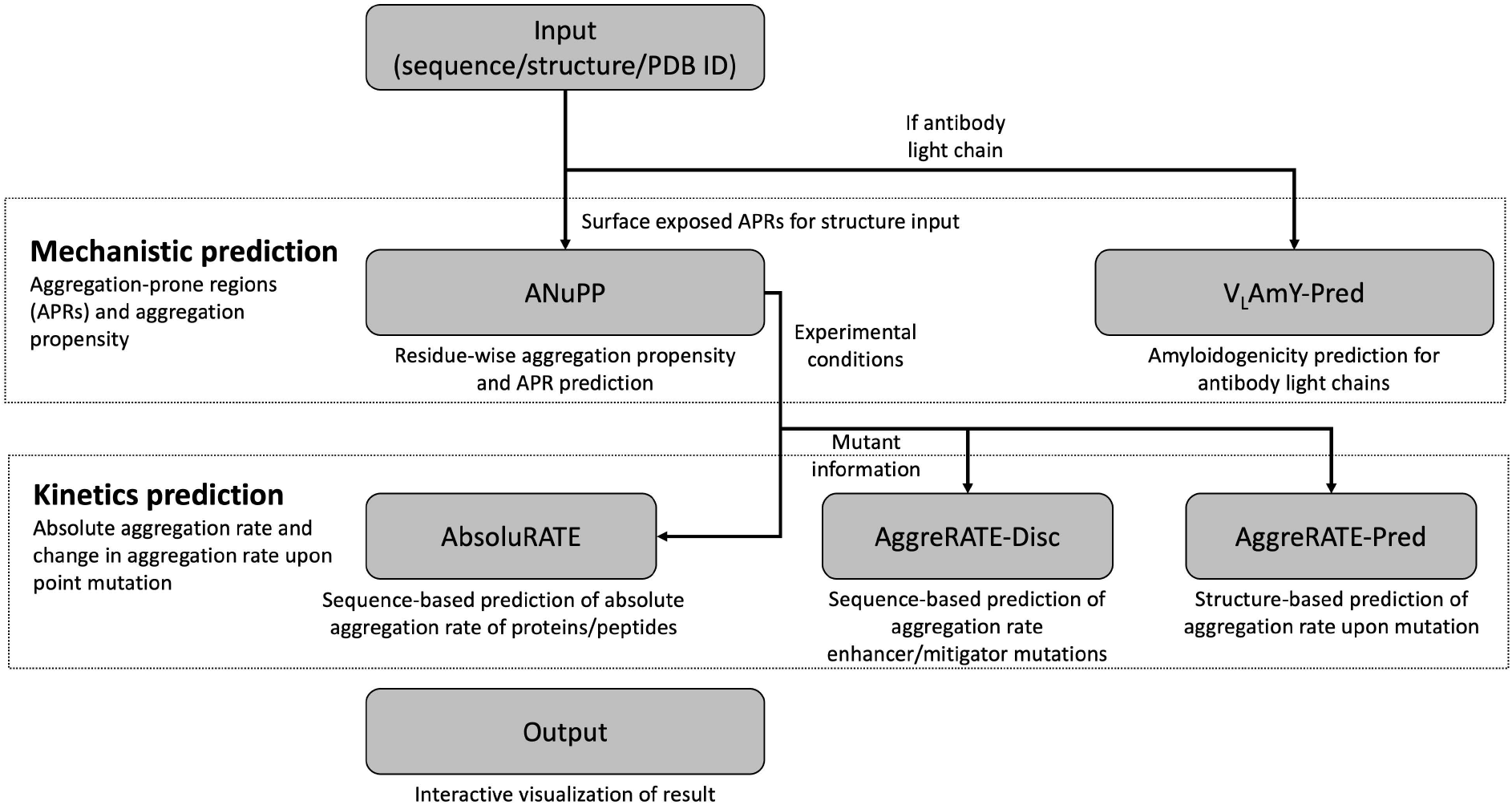
The workflow of the Amylo-Pipe.

## 2. Implementation

Amylo-Pipe is an open-source, automated pipeline that assembles complementary methods to address distinct levels of the aggregation problem. The methods implemented in Amylo-Pipe are discussed in detail below:

### 2.1. Mechanistic prediction of aggregation

The mechanistic prediction methods for aggregation focus on intrinsic aggregation potential and APR identification. These methods are intended to identify sequence segments (APRs) likely to nucleate aggregation based on aggregation propensity scale and, when structural information is available, to prioritize segments that are more plausibly accessible for intermolecular self-association ^14,33^.

#### 2.1.1. Aggregation Nucleation Prediction in Peptides and Proteins (ANuPP)

ANuPP is a sequence-based ensemble-classifier that uses atomic-level features to predict aggregation-prone regions, which has shown a prediction accuracy of 83% on a blind test dataset of 142 peptides and 77% accuracy on 10-fold cross-validation ^19^. ANuPP is used in Amylo-Pipe as the core APR and aggregation propensity predictor. A later benchmarking study across multiple diverse datasets reported that ANuPP showed strong and consistent performance relative to other available APR predictors, making it a reasonable choice as the default mechanistic module in an integrated server ^34^. Surface exposure of the APRs is one of the key aspects to initiate the aggregation process ^21,24,35^. Therefore, Amylo-Pipe extends ANuPP by enabling structural visualisation of predicted APRs, allowing users to assess their surface accessibility in three dimensions. This is a meaningful addition because the APRs buried in the protein core are less likely to contribute towards aggregation than the solvent exposed ones, unless they become transiently or constitutively exposed upon protein misfolding or denaturation.

#### 2.1.2. Prediction of amyloidogenic immunoglobulin (V_**L**_AmY-Pred)

The sequence-based predictor “V_L_AmY-Pred” addresses a more specific but highly relevant problem of predicting the amyloidogenicity of immunoglobulin light chains. Light-chain amyloidosis presents a comparatively well-defined sequence problem with structural context for aggregation prediction, since immunoglobulin framework and complementarity-determining regions provide interpretable contexts for surface exposure, charge balance, and local aggregation risk. The original V_L_AmY-Pred study calculates aggregation-related features (e.g. charge, hydrophobicity) for framework regions (FRs) and complementarity-determining regions (CDRs) to predict the amyloidogenic nature of antibody light chains. V_L_AmY-Pred has shown prediction accuracy of 81.9% on the training dataset (1645 data points) and 71% accuracy on the blind dataset (183 data points) ^30^. Inclusion of this module makes Amylo-Pipe particularly useful for antibody-related aggregation studies and early developability screening, and may also potentially help with diagnosis of patients suffering from diseases caused by light chain amyloidosis ^30^.

### 2.2. Prediction of aggregation kinetics

Current mechanistic predictors identify APRs and rank them based on aggregation propensity, which has been used as a proxy for *in vivo*/*in vitro* change in aggregation capability of protein(s) ^29,36^. However, the time scale of aggregate accumulation depends on additional determinants and may not track simple propensity scores. The aggregation kinetics prediction tools estimate the experimental rate of aggregation (apparent aggregation rates or k_app_) of a known amyloidogenic protein or peptide under defined environmental conditions. We observed on a large-scale protein and peptide dataset that experimental aggregation rate shows a poor correlation with the aggregation propensity ^25^. We have previously developed several aggregation kinetics prediction methods that demonstrated improved performance relative to existing approaches in benchmarking studies ^14^. To complement mechanistic predictions, Amylo-Pipe incorporates dedicated kinetic models that estimate aggregation rates under defined experimental conditions.

#### 2.2.1. Absolute aggregation rate (AbsoluRATE)

The AbsoluRATE method is designed to predict absolute aggregation rates of proteins and peptides by combining intrinsic sequence features with experimental conditions ^25^. The method is developed on a limited but unbiased dataset of 82 unique proteins and obtained a correlation coefficient of 0.72 with mean absolute error (MAE) of 0.91 using leave-one-out cross-validation (LOOCV). The method incorporates temperature, pH, ionic strength, and protein concentration as inputs and performs best under near-physiological conditions.

#### 2.2.2. Aggregation rate prediction upon point mutation (AggreRATE-Disc/AggreRATE-Pred)

Amylo-Pipe also includes mutation-oriented kinetics predictors to estimate whether a point mutation enhances or mitigates aggregation rate and, where applicable, to quantify the expected change. This functionality is valuable for protein engineering as minimal alteration in a protein leading to reduced aggregation often preserves function, expression, and structural integrity. We have included two methods to predict the change in aggregation rate upon point mutation. The first method “AggreRATE-Disc” is a sequence-based classification method to predict aggregation rate enhancer and mitigator mutation ^31^. The machine learning-based method is trained on 220 point mutations and obtained a prediction accuracy of ∼82% using LOOCV. In contrast, AggreRATE-Pred is a structure-informed regression-based mathematical model that obtained LOOCV correlation coefficient of ∼0.73 and MAE of 0.42 on 183 point mutations ^32^. The inclusion of mutation-effect models is also complemented with computational gatekeeper scanning (D, E, R, K, P) in Amylo-Pipe to provide computational protein design support.

## 3. Web Server Interface and Workflow

Amylo-Pipe presents the user interface in modular form (**Fig. 2A**), allowing users to perform mechanistic prediction (**Fig. 2B**) alone or extend the analysis to kinetics prediction when experimental-condition inputs are available (**Fig. 2C**). Amylo-Pipe is implemented using the Django web framework and is accessible through a web-based interface. Sequence or structure input is required for mechanistic analysis, and ANuPP output is returned by default; V_L_AmY-Pred is available for antibody light-chain inputs (**Fig. 2B**). When a structure is supplied, the server further annotates APRs on the structure, helping users distinguish buried from exposed aggregation-prone segments. For kinetics prediction, users may optionally provide a Position-Specific Scoring Matrix (PSSM) file to reduce runtime; otherwise, the server generates it automatically. The experimental conditions are required for the kinetics prediction. To facilitate ease of use, commonly used physiological conditions are provided by default (**Fig. 2C**). After job submission, users can track their job status from the “Jobs” option at the main menu (**Fig. 2D**). The prediction outcome from ANuPP (**Fig. 2E, Fig. 2F**), V_L_AmY-Pred (**Fig. 2G**), AbsoluRATE (**Fig. 2H**), AggreRATE-Disc (**Fig. 2I**), AggreRATE-Pred (**Fig. 2J**) are presented in the result page in modular form. For the structure input, we recommend providing a clean PDB file (avoid non-consecutive or non-standard residue numbering for proteins, especially antibodies) with a single chain to avoid errors in the server. A detailed tutorial is available on the web server at https://web.iitm.ac.in/bioinfo2/amylopipe/help.

**Figure 2.**
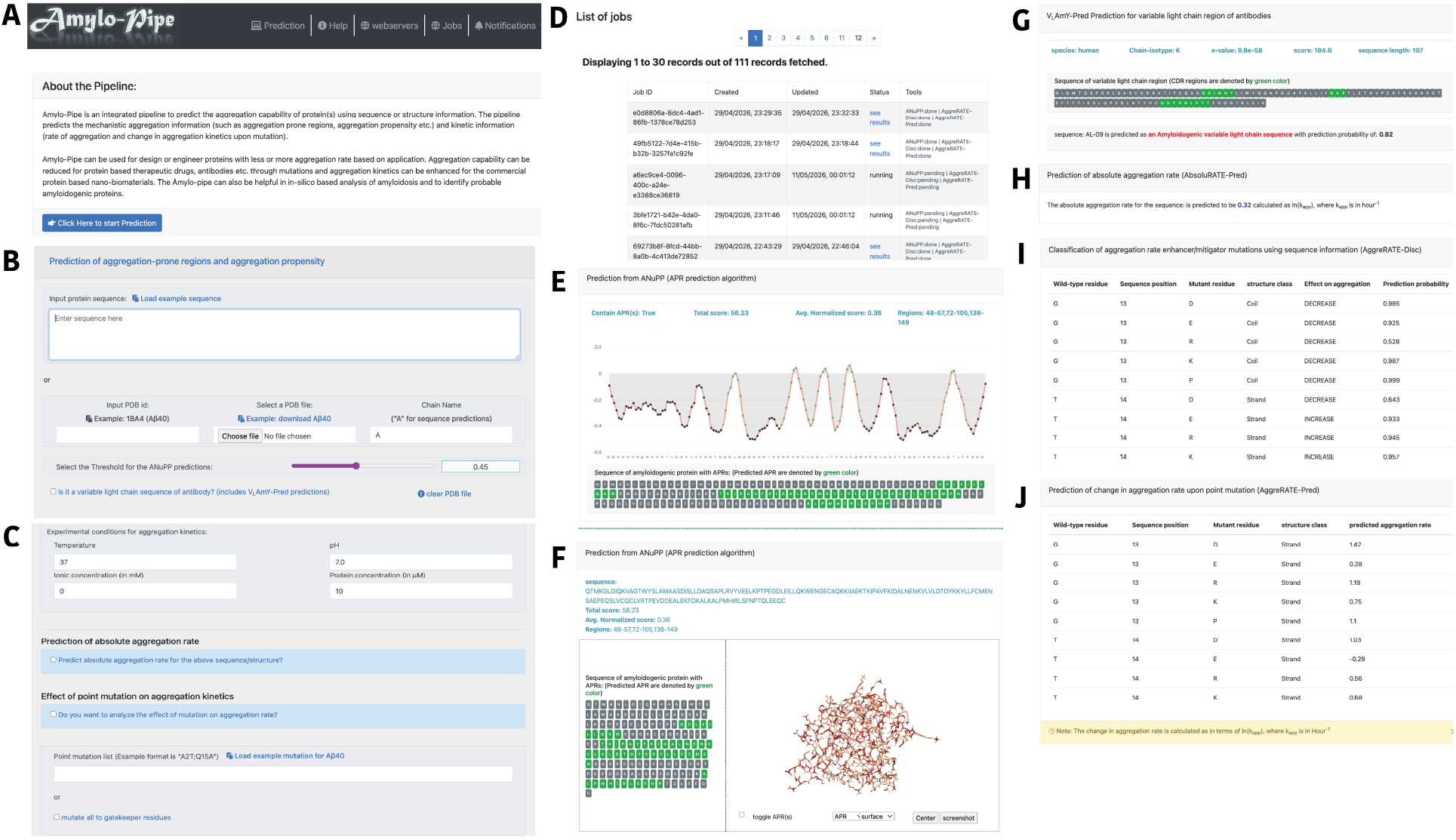
Overview of the Amylo-Pipe webserver. **(A)** The home page of Amylo-Pipe webserver. The prediction page in the server allows users to **(B)** perform mechanistic prediction or **(C)** extend the analysis to kinetics prediction when experimental-condition inputs are available. **(D)** Users can track job status in the Jobs page. It is important to take note of the Job ID if the calculations are expected to take a long time. The result page is modular which presents the results only for the selected models. The sample result page is shown for **(E)** ANuPP (when input is sequence), **(F)** ANuPP (when input is structure), **(G)** V_L_AmY-Pred, **(H)** AbsoluRATE, **(I)** AggreRATE-Disc, **(J)** AggreRATE-Pred.

## 4. Case studies

### 4.1 AcP protein and mutants for analyzing comprehensive aggregation behaviour

We selected the protein with the largest number of experimentally characterized mutations and corresponding aggregation-rate measurements in the CPAD 2.0 database. Acylphosphatase-2 (AcP) comprises 58 single-point mutants evaluated under similar physiological conditions (pH 5.5, 25°C, and protein concentration of 0.4 mg/mL) across four independent studies ^27,37–39^. Amylo-Pipe was applied further to assess the aggregation capability of AcP protein and its variants using structure information. Within the Amylo-pipe framework, ANuPP identified two APRs (residue 6-24 and 32-41) using the default threshold of 0.45 (**Fig. 3A**). Experimental studies have reported six APRs in AcP (residues 16–31, 19–24, 35–54, 87–99, 87–98, and 91–96), which can be clustered into 3 distinct APRs 19–24, 35–54 and 91–96 ^12^. After accounting for overlapping regions, ANuPP successfully identified two of the three experimentally supported non-overlapping APRs (**Fig. 3B**). The C-terminal APR (residues 91–96) was not detected, possibly because of its proximity to the protein terminus; however, the residue-wise aggregation scores showed a marked increase toward the C-terminal region (**Fig. 3A**). The absolute aggregation rate [ln(Kapp), where Kapp was in per hour] for AcP protein was predicted to be -0.05 by AbsoluRATE, whereas the experimental rate was 0.3348 ^27^. To evaluate mutation-level predictions, experimentally measured changes in aggregation rates [Δln(Kapp)] were used to classify variants as aggregation rate enhancer or mitigator, and were compared against AggreRATE-Disc predictions. Overall, 47 of the 58 mutations were correctly classified, corresponding to an accuracy of 81.03%, sensitivity of 83.3%, and specificity of 78.6% (**Fig. 3C, Table 1**). Furthermore, quantitative changes in aggregation rates were compared with AggreRATE-Pred predictions generated using the AcP structure (PDB ID: 1APS). Predicted and experimental values showed strong agreement, with Pearson and Spearman correlation coefficients of 0.73 and 0.59, respectively, and a mean absolute error (MAE) of 0.52 (**Fig. 3D, Table 1**). Notably, the predicted secondary-structure class may differ between AggreRATE-Disc and AggreRATE-Pred because of differences in the input data (sequence versus structure) and the methods used to assign secondary structure.

**Table 1.** Prediction of change in aggregation rate for 58 mutants of Acylphosphatase-2 using Amylo-Pipe platform.

| Mutation information |  |  | AggreRATE-Pred |  | AggreRATE-Disc |  |  | Experimental |  |
| --- | --- | --- | --- | --- | --- | --- | --- | --- | --- |
| WT residue | Position | Mutant residue | Structure class | Predicted ln(Kapp) | Structure class | Predicted change in ln(Kapp) | Prediction probability | delta ln(Kapp) | Change in delta ln(Kapp) |
| A | 30 | G | Helix | 1.48 | Helix | INCREASE | 0.93 | 1.98 | INCREASE |
| E | 29 | D | Helix | -0.52 | Helix | INCREASE | 0.943 | 0.49 | INCREASE |
| E | 29 | K | Helix | -1.08 | Helix | INCREASE | 0.912 | -0.62 | DECREASE |
| E | 29 | Q | Helix | -0.75 | Helix | INCREASE | 0.811 | -0.41 | DECREASE |
| E | 29 | R | Helix | -1.31 | Helix | INCREASE | 0.867 | -1.54 | DECREASE |
| E | 55 | Q | Helix | -0.14 | Helix | INCREASE | 0.833 | 0.13 | INCREASE |
| E | 83 | D | Strand | 0.01 | Strand | DECREASE | 0.634 | -0.08 | DECREASE |
| E | 90 | H | Coil | -1.15 | Coil | DECREASE | 0.792 | -2.52 | DECREASE |
| F | 22 | L | Coil | -1.06 | Helix | DECREASE | 0.682 | -0.32 | DECREASE |
| F | 94 | L | Strand | 0.3 | Coil | DECREASE | 0.963 | -0.09 | DECREASE |
| G | 15 | A | Coil | 0.27 | Coil | DECREASE | 0.865 | 0.05 | INCREASE |
| G | 19 | A | Coil | 0.07 | Coil | DECREASE | 0.699 | 0.36 | INCREASE |
| G | 34 | A | Coil | 0.73 | Coil | INCREASE | 0.869 | -0.2 | DECREASE |
| G | 45 | A | Coil | 0.31 | Coil | INCREASE | 0.662 | 0.22 | INCREASE |
| G | 45 | E | Coil | 0.69 | Coil | DECREASE | 0.56 | 1.09 | INCREASE |
| G | 45 | R | Coil | 0.5 | Coil | INCREASE | 0.896 | 0.51 | INCREASE |
| G | 45 | S | Coil | 0.31 | Coil | INCREASE | 0.92 | 1.08 | INCREASE |
| G | 53 | A | Strand | 0.42 | Coil | INCREASE | 0.549 | 1.19 | INCREASE |
| G | 69 | A | Coil | 0.4 | Coil | INCREASE | 0.938 | 0.71 | INCREASE |
| I | 33 | V | Helix | 0.23 | Helix | INCREASE | 0.825 | 0.05 | INCREASE |
| I | 75 | V | Coil | -0.31 | Strand | DECREASE | 0.908 | -0.11 | DECREASE |
| I | 86 | V | Strand | -1 | Coil | DECREASE | 0.72 | 0.02 | INCREASE |
| K | 67 | A | Strand | 0.17 | Helix | INCREASE | 0.989 | 0.01 | INCREASE |
| K | 88 | N | Coil | -0.52 | Coil | DECREASE | 0.995 | -0.24 | DECREASE |
| K | 88 | Q | Coil | -0.57 | Coil | DECREASE | 0.562 | -0.22 | DECREASE |
| L | 65 | V | Coil | -0.41 | Helix | INCREASE | 0.755 | 0.02 | INCREASE |
| L | 89 | A | Coil | -1.4 | Coil | DECREASE | 1 | -1.79 | DECREASE |
| M | 61 | A | Helix | 0.89 | Helix | INCREASE | 0.528 | 0.04 | INCREASE |
| P | 54 | A | Strand | 0.06 | Coil | INCREASE | 0.854 | 0.16 | INCREASE |
| P | 71 | A | Coil | -0.48 | Coil | INCREASE | 0.991 | 0.03 | INCREASE |
| R | 23 | Q | Coil | -0.82 | Helix | INCREASE | 0.927 | 0.67 | INCREASE |
| R | 97 | E | Strand | 0.07 | Strand | INCREASE | 0.728 | 0.41 | INCREASE |
| R | 97 | Q | Strand | 0.24 | Strand | INCREASE | 0.98 | 2.73308E-05 | INCREASE |
| S | 21 | R | Coil | -1.61 | Coil | DECREASE | 0.972 | -2.06 | DECREASE |
| S | 5 | T | Coil | -0.44 | Coil | DECREASE | 0.902 | -0.15 | DECREASE |
| S | 8 | H | Strand | 0.81 | Strand | INCREASE | 0.821 | 0.09 | INCREASE |
| S | 87 | T | Coil | -0.28 | Coil | INCREASE | 0.889 | 0.59 | INCREASE |
| S | 92 | H | Coil | -0.21 | Coil | DECREASE | 0.597 | -0.34 | DECREASE |
| S | 92 | T | Coil | 0.58 | Coil | INCREASE | 0.678 | 0.88 | INCREASE |
| T | 42 | A | Coil | -0.58 | Coil | DECREASE | 0.723 | -0.1 | DECREASE |
| T | 78 | S | Strand | 0.91 | Strand | INCREASE | 0.966 | -0.16 | DECREASE |
| V | 13 | A | Strand | 0.55 | Strand | DECREASE | 0.788 | -0.13 | DECREASE |
| V | 17 | A | Coil | -0.73 | Strand | DECREASE | 0.885 | -0.26 | DECREASE |
| V | 20 | A | Coil | -1.01 | Coil | DECREASE | 1 | -1.26 | DECREASE |
| V | 36 | A | Strand | 0.58 | Strand | DECREASE | 0.847 | -0.09 | DECREASE |
| V | 39 | A | Strand | -0.52 | Strand | DECREASE | 0.865 | -0.23 | DECREASE |
| V | 47 | A | Strand | -0.01 | Strand | DECREASE | 0.908 | -0.04 | DECREASE |
| V | 51 | A | Strand | 1.06 | Strand | DECREASE | 0.91 | -0.1 | DECREASE |
| V | 9 | A | Strand | 1.14 | Strand | DECREASE | 0.517 | 0.13 | INCREASE |
| W | 64 | F | Strand | 0.21 | Helix | INCREASE | 0.96 | -0.14 | DECREASE |
| Y | 11 | F | Strand | -0.13 | Strand | INCREASE | 0.592 | 0.03 | INCREASE |
| Y | 25 | A | Helix | -0.04 | Helix | DECREASE | 0.986 | -0.52 | DECREASE |
| Y | 91 | Q | Coil | -2.16 | Strand | DECREASE | 0.635 | -3.29 | DECREASE |
| Y | 98 | Q | Coil | -1.41 | Coil | DECREASE | 0.824 | -2.32 | DECREASE |
| K | 44 | E | Coil | 0.68 | Coil | INCREASE | 0.722 | 0.99 | DECREASE |
| Q | 52 | E | Strand | -0.27 | Strand | INCREASE | 0.878 | 0.6 | DECREASE |
| R | 77 | E | Coil | -0.16 | Strand | INCREASE | 0.892 | 0.82 | DECREASE |
| S | 43 | E | Coil | 0.1 | Coil | INCREASE | 0.688 | 0.65 | DECREASE |

**Figure 3.**
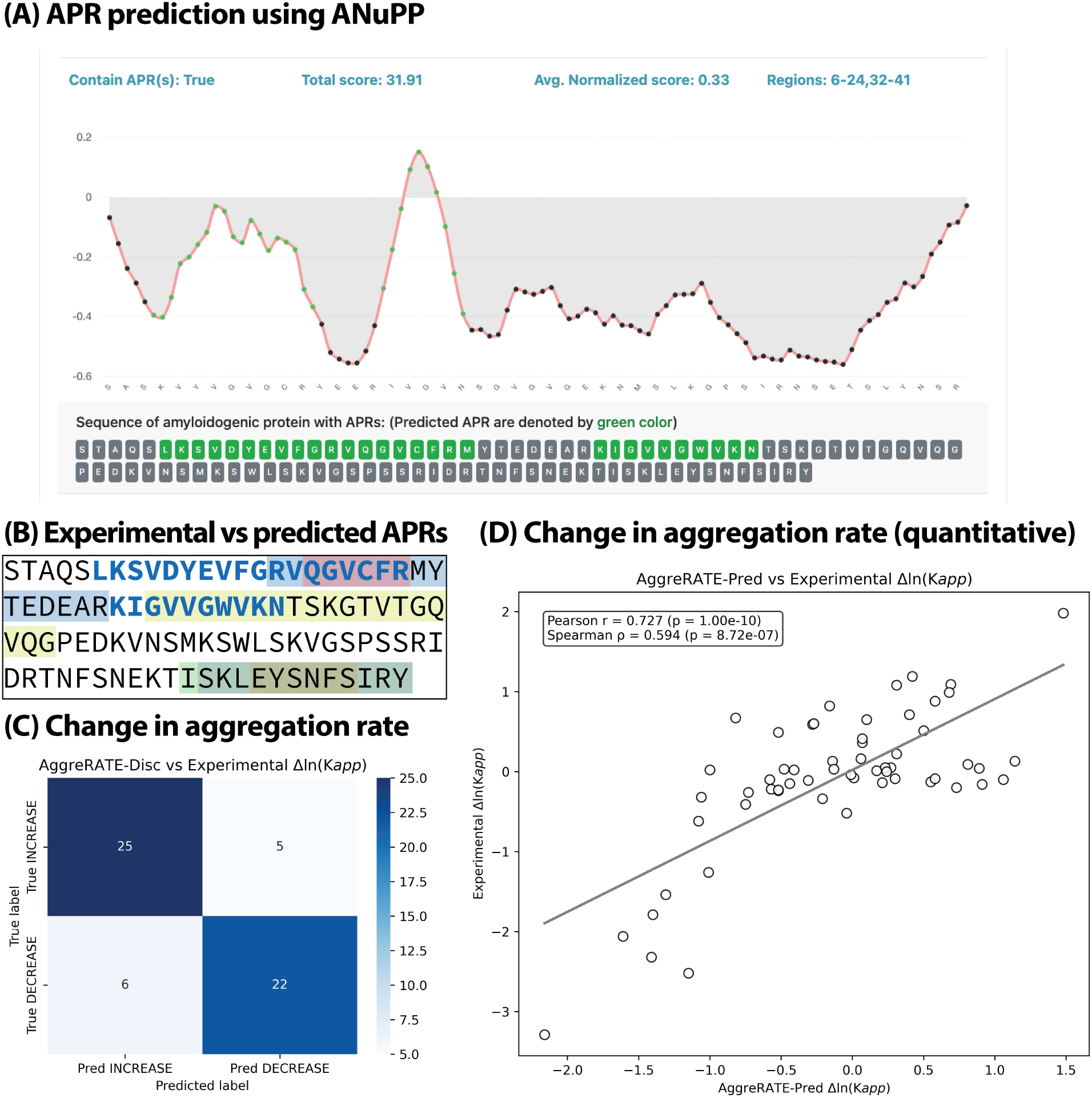
Prediction of aggregation capability of Acylphosphatase-2. **(A)** Residue wise aggregation score and predicted APRs using ANuPP. **(B)** The experimentally identified APRs highlighted in different color boxes, whereas the APR identified by ANuPP are highlighted in blue color and in bold font. **(C)** The increase or mitigation of aggregation prediction by AggreRATE-Disc compared to the experimentally observed change in aggregation rate. The confusion matrix obtained an accuracy of 81.03% (Sensitivity = 83.33%; Specificity = 78.57%). **(D)** The change in aggregation rate predicted by AggreRATE-Pred compared to the experimentally observed change in aggregation rate. A pearson correlation of 0.73, spearman correlation of 0.59 and MAE of 0.5231 was obtained.

### 4.2. Assessment of aggregation behaviour of antibody light chain variable domains

We tested two amyloidogenic antibody light-chain variable domains, AL12 ^40^ and AL103 ^41^ because their experimental aggregation kinetics have been reported. These light chains were not among the training dataset for ANuPP and V_L_AmY-Pred. ANuPP did not identify any APRs in AL12 and detected a single APR in AL103 (residue 28-37; “DISNYLIWYQ”) at default threshold of 0.45, although this region cannot be directly benchmarked at the segment level due to the absence of experimentally mapped APRs for these light chains. However, both AL12 and AL103 were classified as amyloidogenic light-chain sequences by V_L_AmY-Pred, with prediction probabilities of 0.82 and 0.629, respectively. The experimental aggregation rates ln(Kapp) for AL12 and AL103 are −1.91 and −4.40, whereas AbsoluRATE predicted −0.19 and −1.76, respectively, correctly capturing the higher aggregation propensity of AL12 relative to AL103, while underestimating the magnitude of absolute rates.

### 4.3 Assessment of Aggregation Behavior of 0N4R Tau isoform as a blind test dataset

A recent study characterized the aggregation kinetics of the 0N4R tau isoform and 28 disease-associated single-point mutants, providing an independent dataset that was not used to train or validate any of the models implemented in Amylo-Pipe ^42^. This makes 0N4R tau and its variants a suitable external test case to evaluate the practical performance of the web server. Structural modeling of 0N4R tau with AlphaFold2 ^43,44^ yielded low-confidence predictions (low pLDDT), consistent with its intrinsically disordered nature. Consequently, we restricted the analysis to sequence-based predictions and did not apply the structure-dependent AggreRATE-Pred model to this system.

Using ANuPP, we identified two APRs in the 0N4R Tau sequence at residues 238–262 (NIKHVPGGGSVQIVYKPVDLSKVTS) and 330–344 (HGAEIVYKSPVVSGD). To relate these predictions to available experimental evidence, we mapped experimentally validated APRs for canonical tau from CPAD 2.0 onto the 0N4R isoform, yielding six short APRs (length ≤ 20): “GKVQIINKKLDL”, “VQIVYK”, “VQIINK”, “KVQIIN”, “PGGGKVQIVYKPV”, and “PGGGSVQIVYKPVDLSKVTS”. These motifs contain two overlapping core regions, “VQIVYK” and “VQIIN”. At the default ANuPP threshold, “VQIVYK” was recovered. Lowering the threshold to 0.42 introduced an additional APR spanning residues 215–222 (“GKVQIIN”), which captures the experimentally supported “VQIIN” segment.

The experimental aggregation rate [ln(Kapp)] of wild-type 0N4R tau was reported as −0.601, whereas AbsoluRATE predicted a slower apparent rate of −2.34, underestimating the absolute aggregation kinetics. To assess effects of mutations, we classified each tau variant as an aggregation-rate enhancer (6 mutants) or mitigator (22 mutations) relative to wild type and compared the experimental data with AggreRATE-Disc predictions. Across 28 mutants, AggreRATE-Disc correctly classified 20 variants (accuracy 71.43%; sensitivity 81.82%; specificity 33.33%). The lower specificity was observed because the experimental data was biased towards the mutations that increase aggregation rate (**Table 2**).

**Table 2.** Prediction of change in aggregation rate for 28 mutants of 0N4R Tau isoform using Amylo-Pipe platform.

| Experimental detail |  |  |  | AggreRATE-Disc |  |  |
| --- | --- | --- | --- | --- | --- | --- |
| Mutation information | Predicted ln(Kapp) | Predicted change in ln(Kapp) | Structure group | Structure class | Predicted change in ln(Kapp) | Prediction probability |
| A152T | -0.773 | DECREASE | 1 | Coil | DECREASE | 0.711 |
| K257T | -0.724 | DECREASE | 1 | Coil | INCREASE | 0.57 |
| I260V | -0.715 | DECREASE | 2,9 | Coil | DECREASE | 0.982 |
| G272V | -0.501 | INCREASE | 1,9 | Coil | INCREASE | 0.999 |
| G273R | -0.533 | INCREASE | 9,10 | Coil | INCREASE | 0.815 |
| L284R | 0.123 | INCREASE | 9,10 | Coil | DECREASE | 0.996 |
| N296H | 0.022 | INCREASE | 4 | Coil | INCREASE | 0.984 |
| G303V | -1.072 | DECREASE | 1 | Coil | INCREASE | 0.999 |
| G304S | -0.724 | DECREASE | 3 | Coil | INCREASE | 0.96 |
| S305I | -0.355 | INCREASE | 4 | Strand | DECREASE | 0.987 |
| K317M | -0.384 | INCREASE | 4 | Helix | INCREASE | 0.92 |
| P332S | 0.031 | INCREASE | 10,12 | Coil | INCREASE | 0.974 |
| G335V | 0.084 | INCREASE | 1 | Coil | INCREASE | 1 |
| G335S | -0.756 | DECREASE | 2 | Coil | INCREASE | 0.993 |
| Q336R | -0.403 | INCREASE | 13 | Strand | INCREASE | 0.988 |
| V337M | -0.31 | INCREASE | 1,4 | Strand | DECREASE | 0.659 |
| E342V | -0.267 | INCREASE | 1,8 | Coil | INCREASE | 0.999 |
| D348G | 0.158 | INCREASE | 1 | Helix | DECREASE | 0.996 |
| S352L | -0.087 | INCREASE | 4 | Helix | INCREASE | 0.997 |
| V363I | -0.259 | INCREASE | 1 | Coil | INCREASE | 0.558 |
| P364S | -0.368 | INCREASE | 4 | Coil | INCREASE | 0.983 |
| G366R | -0.116 | INCREASE | 4,5 | Coil | INCREASE | 0.815 |
| K369I | -0.02 | INCREASE | 4,7 | Coil | INCREASE | 1 |
| E372G | -0.274 | INCREASE | 11 | Strand | INCREASE | 0.957 |
| G389R | 0.082 | INCREASE | 1 | Coil | INCREASE | 0.777 |
| R406W | -0.239 | INCREASE | 1,2 | Coil | INCREASE | 1 |
| N410H | -0.466 | INCREASE | 1 | Coil | INCREASE | 0.998 |
| T427M | -0.054 | INCREASE | 1 | Helix | INCREASE | 0.934 |
| **WT** | **-0.601** |  | **1** | **-** | **-** | **-** |

The experimental study further grouped mutant-derived fibrils into 13 structural classes based on fibril conformation (**Table 2**), highlighting that many mutants alter fibril structures relative to the wild type. To assess whether our misclassifications were driven by fibril conformation, we examined the distribution of structural classes among correctly and incorrectly predicted variants. Among the eight mutants misclassified by AggreRATE-Disc, four belonged to structure group 1 (fibrils structurally similar to wild type), and the remaining four were scattered across different structural groups without a clear pattern. Likewise, among the 20 correctly classified mutants, nine were assigned to structure group 1. This distribution indicates no systematic dependence of prediction accuracy on fibril conformation in this dataset. Overall, this external test case supports the use of Amylo-Pipe to analyse aggregation behaviour of proteins and their variants, with kinetics models that are useful but clearly not perfect yet. This observation re-emphasizes the need for collection of greater experimental data on aggregation kinetics.

## 5. Potential application of the webserver

Amylo-Pipe offers an integrated framework for assessing aggregation-prone regions and aggregation kinetics, with applications in developability assessment, protein engineering, and disease-focused aggregation studies. It is particularly useful for comparative analysis of protein variants under user-defined experimental conditions. For therapeutic proteins and antibodies, the server can screen candidates for aggregation liability and suggest sequence changes that may reduce aggregation while preserving structure. Gatekeeper scanning and mutation-oriented kinetics models support rational design by pinpointing positions where substitutions are predicted to mitigate aggregation rates. In antibody light chains, combining VLAmY-Pred with APR and kinetics modules enables simultaneous evaluation of amyloidogenicity and local aggregation hot spots. In fundamental research, Amylo-Pipe helps generate hypotheses on how mutations, sequence motifs, or environmental parameters influence aggregation mechanisms and kinetics, guiding the design and interpretation of aggregation kinetic experiments. While not a replacement for real-time stability or shelf-life measurements, its predictions can narrow the experimental search space and provide mechanistic context for studies of protein aggregation.

## 6. Discussion

Amylo-Pipe is designed to overcome a practical limitation of the current ecosystem of aggregation-prediction servers. Most available tools and their web servers provide only mechanistic predictions, focusing on identification of potential APRs in a protein and its overall aggregation propensity ^18,22,24,45–47^. Predictions of aggregation kinetics are not provided in most cases. Amylo-Pipe bridges this gap by integrating mechanistic as well as kinetic predictors of protein and peptide aggregation via a single workflow. The Amylo-Pipe web server is user-friendly and has been optimised to minimise prediction time and provide comprehensive information on the aggregation behaviour of a protein or peptide.

Regarding the choice of predictors included in this workflow, ANuPP was chosen to identify APRs because it improves upon classical residue-level methods by incorporating atomic-level features, and has demonstrated significantly better and more consistent performance across diverse datasets including hexapeptides, proteins with known APRs, homologous proteins, and aggregating antibodies ^34^. A practical strength of ANuPP within Amylo-Pipe is that it extends sequence-based APR prediction by structurally visualizing APRs on protein structure for surface-exposure.

Light chain amyloidosis remains a life-threatening condition with limited therapeutic options. Although most APR prediction algorithms perform reasonably well on globular proteins, they have not been trained or validated on immunoglobulin sequences ^30,34^. In the case of antibody light chains, sequence information can provide crucial detail of exposed hydrophobic regions (CDRs of antibody) and gatekeeping effect for exposed regions (FRs in antibody). These sequence features form the basis of V_L_AmY-Pred, enabling accurate classification of amyloidogenic antibody light chains.

The experimental aggregation kinetics of a protein (k_app_) depend on several other factors, including surface exposure of APRs and environmental conditions such as pH, temperature, ionic strength, protein concentration, agitation stress and solution buffer composition. We previously collected the most comprehensive amyloid database “CPAD 2.0”, which also includes aggregation kinetics of amyloidogenic proteins along with experimental conditions ^12^. Using CPAD 2.0, we have developed robust kinetics prediction methods: AbsoluRATE for absolute aggregation rate prediction ^25^, AggreRATE-Disc for classifying rate-enhancing and rate-mitigating mutations ^31^, and AggreRATE-Pred for quantifying the change in aggregation rate upon point mutation ^32^. As far as we know, there are no additional aggregation kinetics programs available in public domain, thereby precluding the opportunity of an informed selection. Amylo-Pipe also features a gatekeeper mutational scanning capability that enables users to simultaneously mutate all protein positions with gatekeeper residues (R, K, D, E, P) to assist in the design of aggregation-resistant proteins or peptides.

While Amylo-Pipe integrates validated methods, there are also several limitations. First, aggregation prediction remains constrained by the availability and experimental heterogeneity of curated training datasets, particularly for kinetics. Moreover, the output of APR prediction, amyloidogenicity classification, and rate prediction should not be interpreted as interchangeable measures, as these methods are developed independently. Structure-based predictions are generally considered more reliable. For the structure-based analysis, all methods rely on supplied experimental or modeled structure and may not fully capture conformational dynamics, transient exposure, or environmental perturbations.

## 7. Conclusion

Amylo-Pipe is a collection of state-of-the-art mechanistic and kinetics prediction methods related to protein aggregation. It also implements user-friendly features including automated calculation from sequence or structure inputs, structural visualisation of APRs, gatekeeper mutational scanning, and job tracking to minimise wait time. Amylo-Pipe is expected to serve as a valuable resource for *in silico* amyloidogenicity analysis and developability screening of therapeutically relevant proteins and peptides, including antibodies.

## Availability and Implementation

The webserver is open source and accessible at https://web.iitm.ac.in/bioinfo2/amylopipe/.

## Acknowledgements

We thank Bioinformatics Infrastructure facility, Department of Biotechnology and Indian Institute of Technology Madras for computational facilities and Ministry of human resource and development (MHRD) for HTRA scholarship to PR.

